# Leveraging mRNA platform for the development of vaccines against egg allergy

**DOI:** 10.1101/2025.03.14.643193

**Authors:** Xianyu Shao, Changzhen Weng, Lijing Liu, Kun Guo, Zhutao Lu, Lulu Huang, Zhenhua Di, Yixuan Guo, Guorong Di, Renmei Qiao, Jingyi Wang, Yong Yang, Shiyu Sun, Shentian Zhuang, Ang Lin

## Abstract

Food allergy has posed a major global health burden due to the rising prevalence and lack of effective prophylactic strategies. During the past decades, allergen-specific immunotherapy (AIT) has been used as a disease-modifying therapy for food allergic conditions, but due to the long-term treatment duration, poor patient compliance and unexpected adverse reactions, only a minority of patients benefit from AIT therapy. In this proof-of-concept study, using well-established mRNA platform, we developed mRNA vaccine candidates encoding for the major egg white allergen Gal d2 (also known as ovalbumin) and comprehensively evaluated the prophylactic efficacy against anaphylaxis in a Gal d2-induced allergic mouse model. Two vaccine formulations, Gal d2 mRNA vaccine and Gal d2-IL-10 mRNA vaccine, both demonstrated effective ability in preventing the onset of allergic disease, which is largely attributed to the versatilities of mRNA vaccine in eliciting allergen-blocking antibody, shifting Th2 towards Th1 immunity, as well as in generating peripheral tolerance.

---

Allergic diseases, also referred to as type I hypersensitivity reactions, have posed a major global health burden due to the rising prevalence and lack of effective prophylactic strategies. Among various types of allergic conditions, food allergy (FA) represents a critical category and affects about 8% of children and 10% of adults in developed countries ^[1]^. In Chinese population, prevalence of FA has been increasing rapidly and was reported to range from 4.0% to 8.2% according to a meta-analysis ^[2]^. Currently, strict self-avoidance of specific food is commonly used as a strategy by majority of FA patients, but accidental allergen exposure still occurs frequently which requires immediate rescue medication otherwise a life-threatening systemic allergic reaction known as anaphylaxis may appear in some severe cases. During the past decades, allergen-specific immunotherapy (AIT) has been used as a disease-modifying therapy for food allergic conditions including peanut allergy, wheat allergy, etc., which is practiced by exposing the patients to gradually increasing amounts of allergen in controlled dosing to induce tolerance ^[3]^. However, due to the long-term treatment duration, poor patient compliance and unexpected adverse reactions, only a minority of patients benefit from AIT therapy. Therefore, effective prophylactic interventions are largely needed for FA patients.

Allergic disorders are typically characterized by aberrant generation of Th2-polarized immune responses indicated by induction of allergen-specific IgE and Th2-type cytokines (IL-14, IL-5, IL-13, etc.) secreting T cells that collaboratively lead to activation of effector cells such as basophils and mast cells, which play a critical role in mediating and exaggerating allergic reactions ^[4]^. Taking advantages of the principles of immune system, considerable attempts have been made to develop anti-allergy vaccines to prevent FA incidence by inducing allergen-blocking antibodies competing with IgE and modulating or even converting allergen-specific Th2 immunity to Th1 phenotype ^[5]^. Using defined food allergens as immunogens, vaccine modalities based on virus-like particle (VLP) platform, DNA platform, viral-vectored platform and adjuvanted protein-based subunit vaccine platform have been tested and showed promise in mitigating allergic response to food allergens ^[6-9]^. While compared with traditional vaccine types, mRNA vaccine is superior at inducing stronger Th1-type T cell responses and higher magnitude of Ab responses. Moreover, mRNA vaccine formulated using lipid-nanoparticle (LNP) delivery system was reported to induce regulatory T cell (Treg) response contributing to immune tolerance ^[10, 11]^, which provides additional advantage to mRNA technology for anti-allergy vaccine development. In this proof-of-concept study, using well-established mRNA platform ^[12-14]^, we developed mRNA vaccine candidates encoding for the major egg white allergen Gal d2 (also known as ovalbumin) and comprehensively evaluated the prophylactic efficacy against anaphylaxis in a Gal d2-induced allergic mouse model.

Gal d2-mRNA was codon-optimized using a proprietary artificial intelligence-based algorithm and synthesized by in-vitro transcription procedure with N1-methyl-pseudouridine (m1Ψ) incorporated as previously reported ^[12]^, which demonstrated efficient protein expression following transfection into HEK-293T cells (Fig. 1a). Vaccine formulation was prepared by packaging the mRNAs into SM102-containing LNP system using a microfluidics-based procedure. To study the immunogenicity and anti-allergy effect of vaccine, BALB/c mice were administered intramuscularly (i.m.) with three doses of vaccine (1μg or 5μg) at an interval of 7 days. Thereafter, mice were sensitized twice via intraperitoneal (i.p.) administration with Gal d2 followed by intragastric (i.g.) allergen challenge four times consecutively, and anaphylaxis was finally induced through i.p. injection with Gal d2 (Fig. 1b). At day 33 prior to Gal d2 sensitization, mice immunized with Gal d2-mRNA vaccines produced robust levels of Gal d2-specific IgG and two IgG subclasses in a dose-dependent manner (Fig. 1c). Anti-Gal d2 IgE was undetectable therefore excluding the possibility that mRNA vaccine might elicit IgE response (Fig. 1d). In addition, Gal d2-mRNA vaccination elicited strong Th1-type T cells secreting IFN-γ or IL-2 which is quite expected (Fig. 1e-f). As a control group, unvaccinated mice showed no or very limited background levels of Ab or T cell responses (Fig. 1c-f).

**Fig. 1.**
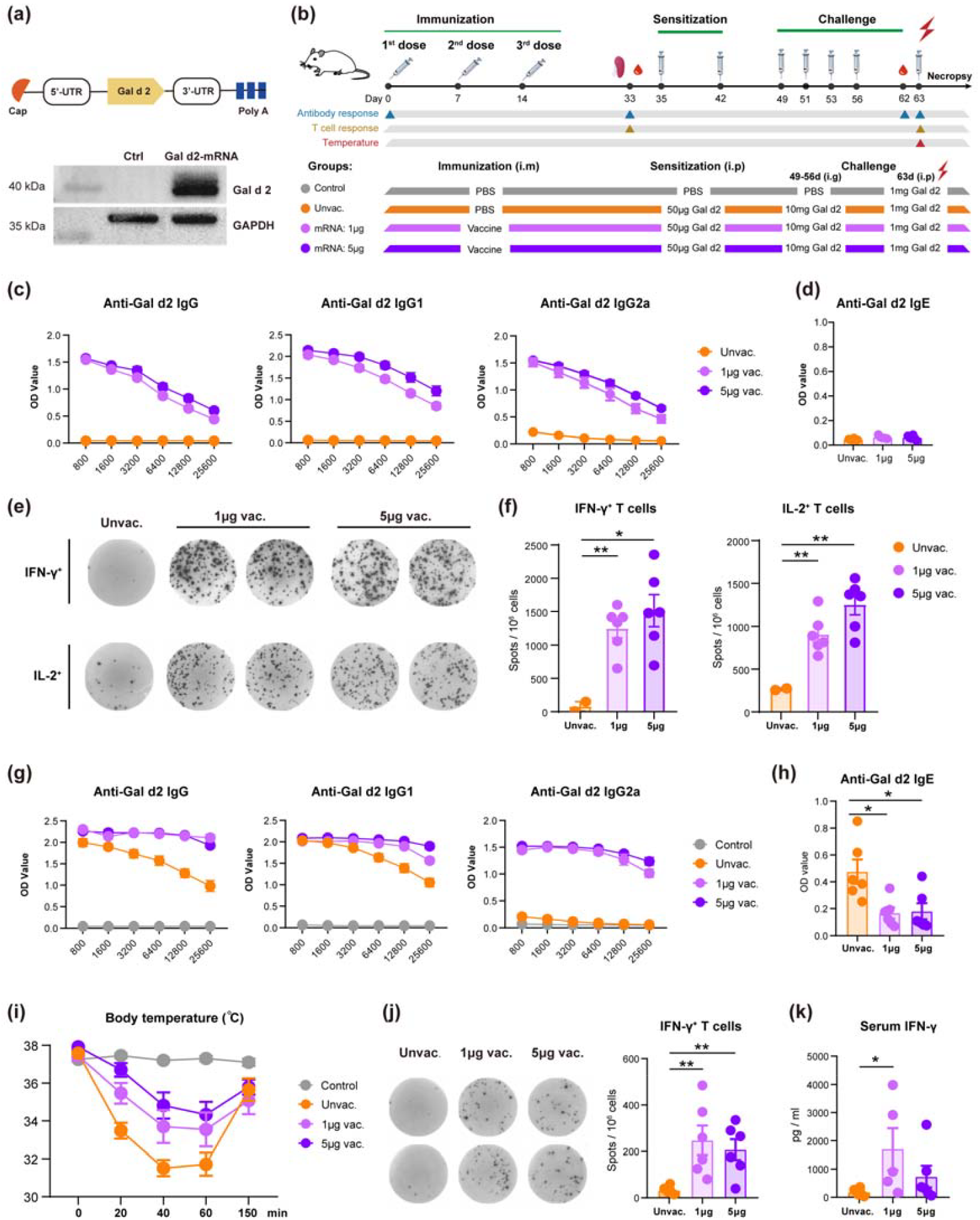
Immunogenicity and anti-allergy efficacy of Gal d2 mRNA vaccine. (a) Construction of Gal d2-mRNA and its translational efficiency in HEK-293T cells was assessed by western blot. (b) Experimental design. BALB/c mice (n=12) were immunized three times at day 0, 7, and 14, followed by Gal d2 sensitization and i.g. challenge consecutively. Final Gal d2 challenge was i.p. administered to induce anaphylaxis. (c-d) Gal d2-specific IgG, IgG1, IgG2, and IgE in mice (n=6) were measured at day 33. (e-f) Frequencies of Gal d2-specific IFN-γ or IL-2-producting T cells in spleens were measured by ELISpot assay. (g-h) At day 62 prior to final allergen challenge, Gal d2-specific IgG, IgG1, IgG2, and IgE in mice (n=6) were measured. (i) Following final i.p. allergen challenge, rectal temperature of mice was monitored. (j) Four hours post the final i.p. allergen challenge, levels of Gal d2-specific IFN-γ or IL-2-producting T cells were quantified (n=6) and serum level of IFN-γ was measured (k). Mann-Whitney U test was used for statistical analysis. *p ≤ 0.05, **p ≤ 0.01.

Following the immunization procedure, animals undergone a series of allergen sensitization and challenge schedule to achieve allergic status to Gal d2. At day 62 prior to the final i.p. injection to induce anaphylaxis, levels of Gal d2-specific IgG and IgE were evaluated. Mice that had been immunized with vaccine maintained high levels of specific IgG, IgG1 and IgG2a, which were even at a higher magnitude than that detected at day 33 (Fig. 1g, 1c). This can be explained by the continuous exposure to antigen during sensitization and challenge stages that boosted memory B cell pool. While interestingly, unvaccinated mice that had undergone allergen sensitization and challenge regimen also produced a robust level of specific IgG that was exclusively composed of IgG1 subclass, however, the Th1-prone antibody IgG2a was barely induced (Fig. 1g). This suggested that Gal d2-allergic mice presented aberrant allergen-specific Th2-type immune signature. Notably, unvaccinated Gal d2-allergic mice showed a significantly higher level of allergen-specific IgE than the two groups of mice that had been vaccinated (Fig. 1h), which demonstrated protective potential of Gal d2-mRNA vaccine since IgE is the key mediator of allergic response. Further, anaphylaxis was induced via i.p. challenge with Gal d2 and rectal temperature as the key indication of symptom was monitored. Mice in the control group (PBS treated) showed steady body temperature upon allergen challenge. However, unvaccinated allergic mice showed a sharp decline in temperature with a maximal drop observed at 40 min, and the temperature gradually recovered to normal state by 150 min (Fig. 1i). In contrast, mice that were immunized with Gal d2-mRNA vaccine showed a noticeably milder temperature drop indicating a desired anti-allergy efficiency conferred by mRNA vaccine, which was more prominently seen in the 5μg dosing group. We also evaluated Gal d2-specific T cell responses 4 hours after the final allergen challenge. Compared with the unvaccinated group, immunized mice showed robust and higher frequencies of IFN-γ-producing T cells (Fig. 1j) and higher levels of serum IFN-γ (Fig. 1k). These collectively suggested that pre-vaccination with Gal d2-mRNA vaccine was able to induce allergen-specific Th1-type T cell immunity that counteracts with the generation of Th2-prone immunity, which may contribute to the manifestations of anti-allergy effect.

Given the above finding showing promising anti-allergy effect of Gal d2-mRNA vaccine, we further modified the vaccine modality by conjugating immune-suppressive cytokine IL-10 encoding mRNA with Gal d2-mRNA through T2A peptide (Fig. 2a). IL-10 is essential not only for the induction of peripheral tolerance to allergen but also plays a critical role in the suppression of IgE response ^[15]^. Upon transfection into HEK-293T cells, the constructed Gal d2-IL-10 mRNA was efficiently translated with a robust level of IL-10 detected in the supernatant (Fig. 2a). We further compared the immunogenicity and anti-allergy efficiency between Gal d2-IL-10 mRNA vaccine and Gal d2 mRNA vaccine according to an identical experimental schedule as shown earlier (Fig. 2b). In terms of antibody response, triple doses of 5μg Gal d2-IL-10 mRNA vaccine induced a comparable level of IgG and IgG2a to Gal d2 mRNA vaccine, while Th2-prone antibody IgG1 was elicited at a lower level by Gal d2-IL-10 mRNA vaccine (Fig. 2c). Again, anti-Gal d2 IgE was not elicited in all vaccine groups (Fig. 2d). Moreover, robust levels of antigen-specific IFN-γ or IL-2-producing T cells were induced by the two vaccine formulations with no difference observed. While noticeably and interestingly, IL-10-secreting T cells were generated in all vaccine groups, which was even more robust in Gal d2 mRNA vaccine group (Fig. 2e-f). These vaccine-elicited IL-10^+^ T cells are likely functional in mitigating allergic response, and the in-depth phenotyping and characterization would merit further investigation. Following sensitization and challenge schedule and prior to the final challenge to induce anaphylaxis, IgG, IgG1 and IgG2a titers were further boosted in all vaccine groups (Fig. 2g), which was in line with the earlier finding (Fig. 1g). Anti-Gal d2 IgE titer was significantly reduced in vaccinated animals but showed no difference between the two vaccine formulations (Fig. 2h). Following final allergen challenge, rectal temperature was monitored, and all vaccinated mice were protected from severe temperature decline during anaphylaxis when compared with the unvaccinated group. On note, Gal d2-IL-10 mRNA vaccine showed more effective ability in alleviating allergic response than the Gal d2 mRNA vaccine, although not significantly (Fig. 2i). Four hours after allergen challenge, we evaluated the frequencies of activated basophils (IgE^+^CD200R3^+^CD63^+^) in spleens that are major effector cells mediating allergic reaction. This specific cell population was barely detected in the control PBS-treated mice but was present at a high level in the unvaccinated allergic mice. In contrast, vaccinated mice showed relatively lower frequencies of activated basophils with no clear difference between the two vaccine formulations (Fig. 2j).

**Fig. 2.**
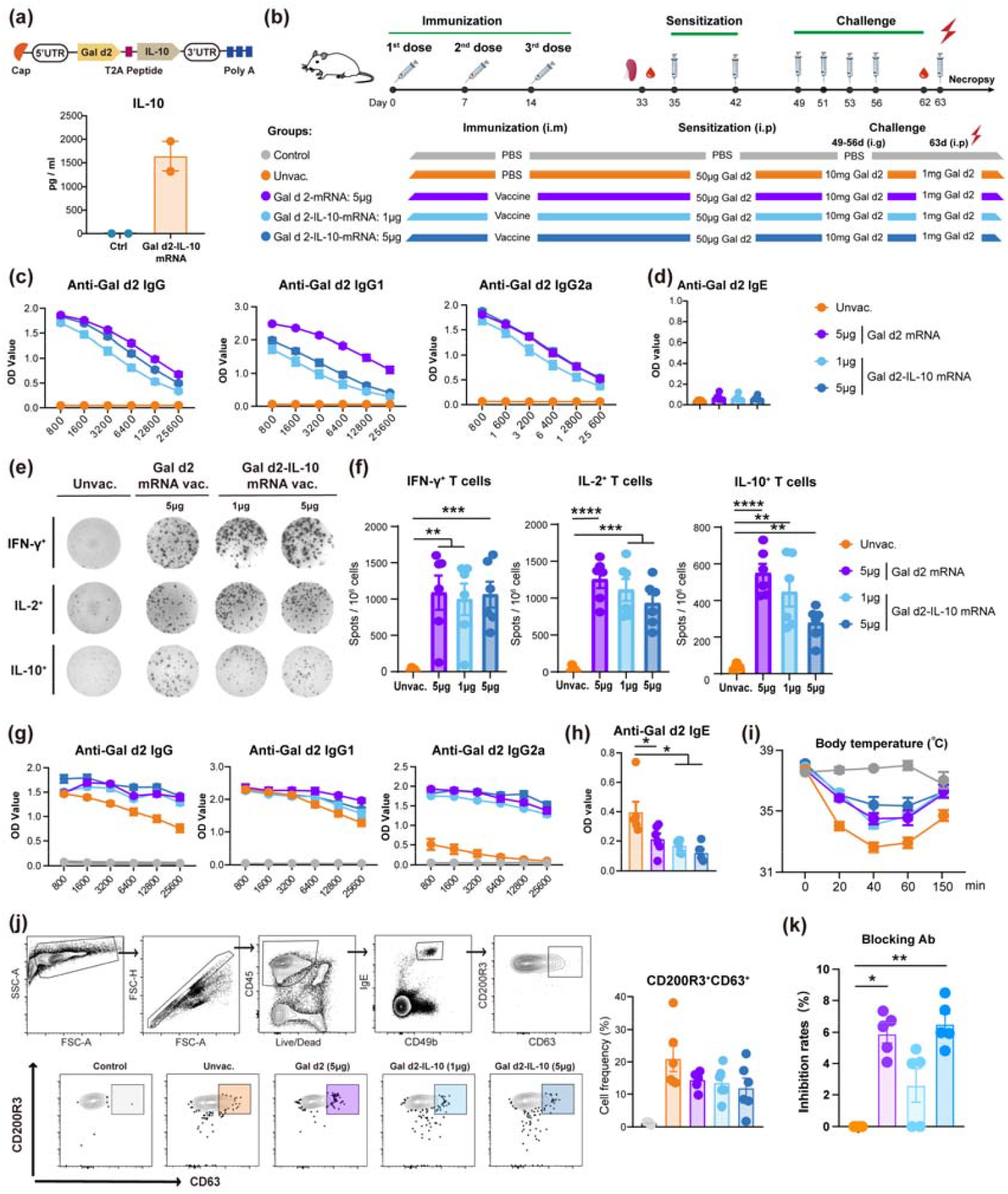
Immunogenicity and anti-allergy efficacy of Gal d2-IL-10 mRNA vaccine. (a) Construction of Gal d2-IL-10 mRNA (upper panel). Expression of IL-10 in the culture supernatant was determined by ELISA following mRNA transfection into HEK-293T cells (lower panel). (b) Experimental design. BALB/c mice (n=12 each group) were immunized three times at day 0, 7, and 14, followed by Gal d2 sensitization and i.g. challenge consecutively. Final Gal d2 challenge was i.p. administered to induce anaphylaxis. (c-d) Gal d2-specific IgG, IgG1, IgG2, and IgE in mice (n=6 each group) were measured at day 33. (e-f) Frequencies of Gal d2-specific IFN-γ, IL-2, or IL-10-producting T cells in spleens were measured by ELISpot assay. (g-h) At day 62 prior to final allergen challenge, Gal d2-specific IgG, IgG1, IgG2, and IgE in mice (n=6) were measured. (i) Following final i.p. allergen challenge, rectal temperature of mice was monitored. (j) Four hours post the final i.p. allergen challenge, frequencies of activated basophils were measured by FACS. (k) Allergen-blocking capacity of vaccine-induced antibodies was measured (n=5 each group). Inhibition rate is shown. One-way ANOVA test was used for statistical analysis. *p ≤ 0.05, **p ≤ 0.01, ***p ≤ 0.001, ****p ≤ 0.0001.

Collectively, Gal d2 mRNA and the optimized Gal d2-IL-10 mRNA vaccines demonstrated comparable ability in preventing the onset of allergic disease. With regards to the potential mechanisms, induction of strong Th1-type immunity counteracting with Th2-type allergic immunity (Fig. 1e, 2e) and the generation of regulatory IL-10^+^ T cells (Fig. 2e) may play a critical role. In addition, the mRNA vaccine-elicited IgG could function as blocking antibody competing with IgE for binding to the allergen. This was supported by competitive ELISA experiments showing that vaccine-elicited antibodies were able to block IgE-allergen interaction (Fig. 2k). In this brief study, we provided preliminary evidence showing that mRNA platform is unique and holds promise for the development of anti-allergy vaccines. This is largely attributed to the versatilities of mRNA vaccine in eliciting allergen-blocking antibody, shifting Th2 towards Th1 immunity, as well as in generating peripheral tolerance. However, there are some weaknesses of this study that await to be further addressed, such as therapeutic potential of mRNA vaccine in treating existing food allergy, influence of routes of administration on vaccine efficacy, mechanisms behind the induction of regulatory T cells by mRNA vaccine, etc. In-depth investigations would be required to gain deep mechanistic insights into the preventive efficacy of mRNA vaccine that will provide guidance for rational anti-allergy vaccine development.

## Acknowledgements

The authors thank the Targeted Discovery Center of China Pharmaceutical University for instrumental support on this study. This work was supported by National Natural Science Foundation of China (Grant nr. 32471004, 32200764 to A.L.; 32301241 to S.Z.), Innovation Capacity Building Initiative Funding (Grant nr. BM2023002 to Y.Y.). X.S. and K.G. were also supported by the Postgraduate Research & Practice Innovation Program of Jiangsu Province.

## Author contributions

A.L., S.Z., S.S. and Y.Y. designed research. X.S., C.W., L.L., K.G., Z.L., L.H., Z.D., Y.G., G.D., R.Q., J.W. performed the experiments; X.S., C.W., L.L., K.G. and A.L. analyzed the data; X.S., C.W., S.S., S.Z. and A.L. discussed the data. X.S., L.L. and A.L. wrote the manuscript. All authors provided critical review of the manuscript.

## Appendix A. Supplementary materials

Detailed description of the materials and methods used in this study can be found in the supplementary materials.

