## Supplementary materials for "Leveraging mRNA platform for the development of vaccines against egg allergy"

**Materials and Methods**

**Ethics and animals**

BALB/c mice (female, 6-week-old) were purchased from GemPharmatech Co., Ltd. (Nanjing, China) and randomly allocated to different groups. Mice were kept in specific pathogen-free (SPF) condition at the Center for New Drug Safety Evaluation and Research at China Pharmaceutical University. All animal experiments were performed in accordance with the Guidelines for the Care and Use of Laboratory Animals and the Ethical Committee of China Pharmaceutical University using protocols approved by the Institutional Animal Care and Use Committee of China Pharmaceutical University (approval number: VR-B2301P090).

**mRNA vaccine preparation**

mRNAs encoding for Gal d2 or Gal d2-IL-10 were codon-optimized and synthesized by T7 polymerase-mediated in vitro transcription (IVT) using a linearized DNA template (pUC57-GW-Kan) containing 5’ untranslated regions (UTRs), 3’UTRs and a 120 nt poly-A tail. During the IVT procedure, N1-Methyl-pseudouridine (Synthgene) was used to modify mRNAs and mRNAs were capped using Cap 1 Analogue Reagent (Synthgene). Subsequently, IVT products were purified using Monarch RNA purification columns (NEW ENGLAND BioLabs Inc. MA, USA) and resuspended in TE buffer. The lipid components were dissolved in ethanol at molar ratios of 50:10:38.5:1.5 (SM102: DSPC: cholesterol: DMG-PEG2000, purchased from SINPOEC). mRNAs were dissolved in 10 mM citrate buffer (pH 4.0) and then encapsulated into lipid nanoparticles (LNPs) at a volume ratio of 3: 1 using a microfluidic-based device (INano^TM^L from Micro&Nano Biologics) at a flow rate of 12 mL/min. Formulations were diluted with PBS and ultrafiltrated using 50-kDa Amicon ultracentrifugal filters. Vaccine formulations were characterized by NanoBrook Omni ZetaPlus (Brookhaven Instruments) for particle diameter, polymer dispersity index and zeta potentials.

**Cell culture**

HEK-293T cells were purchased from Cell Resource Center, Shanghai Institutes for Biological Sciences, Chinese Academy of Sciences and cultured in DMEM (BIOIND, Israel) supplemented with 10% fetal bovine serum (FBS, BIOIND, Israel) and 1% penicillin-streptomycin (NCM Biotech, China). All cells were maintained at 37 ℃ and in a 5% CO_2_ atmosphere.

**Immunization and sampling**

To evaluate the immunogenicity and preventive effect of vaccine, three doses of Gal d2 mRNA or Gal d2-IL-10 mRNA vaccines were i.m. administered to BALB/c mice (start with n=12 per group) at an interval of 7 days. Sera samples (n=6 per group) were collected longitudinally at indicated time points for analyses of antibody responses. Following immunization schedule, mice from indicated groups were sensitized by i.p. injection of 50 μg of Gal d2 (Sigma Aldrich) mixed with 2 mg of alum adjuvant (Thermo Scientific) per dose at day 35 and 42, followed by i.g. challenge with 10 mg of OVA at day 49, 51, 53 and 56. One week post the final i.g. challenge, mice were i.p. challenged with 1 mg of Gal d2 to induce anaphylaxis. Rectal temperature at indicated time points was recorded with a rectal thermometer. Four hours post the final i.p. challenge, animals were sacrificed, and spleens were processed to obtain single cell suspension. Overview of the experimental schedule is displayed in Figure 1b and Figure 2b.

**Preparation of single cell suspension**

Spleen tissues were grounded gently and filtered through a 70-μm sterile cell strainer. Cells were resuspended in PBS and centrifuged at 400 g for 10 min. To lyse red blood cells (RBC), cell pellets were resuspended with RBC lysis buffer (Solarbio) for 5 min at 4 °C. Following this, 1× PBS was added to terminate the lysis procedure and cells were then washed at 400 g for 10 min to obtain splenic MNCs. Thereafter, cells were re-suspended in RPMI-1640 medium containing 10% FBS (BIOIND) and 1% penicillin-streptomycin (NCM Biotech) for subsequent in-vitro experiments.

**Western blot assay**

HEK-293T cells were transfected with mRNAs using jetMESSENGER® transfection reagent according to the instruction. Upon 24 hours of incubation, cells were collected and lysed, followed by centrifugation (12,000 rpm, 15 min, 4 ℃) for protein extraction. Protein concentration was determined by BCA Protein Quantification Kit-BOX 2 (Vazyme). 20 μg of proteins were loaded onto a 10% SDS-polyacrylamide gel for SDS-PAGE electrophoresis. Proteins were then transferred onto PVDF membranes and were blocked with 5% milk for 2 hours at room temperature (RT). The membrane was then washed with TBS containing 0.075% Tween-20 (TBST) for 3 times and incubated with 1: 10000 diluted anti-Gal d2 antibody (Proteintech, CloneNo.1D3D5) at 4 ℃ overnight. After washing steps, the membrane was incubated with 1: 10000 diluted HRP-conjugated goat anti-mouse IgG (Fdbio science) for 2 h. The membrane was washed, and spots were visualized using Amersham ECL Prime Western Blotting Detection Reagent (GE Healthcare).

**Measurement of Gal d2-specific antibody titer**

96-well plates (Greiner Bio-One) were pre-coated with Gal d2 antigen (100 µg/mL) at 4 ℃ overnight. The plates were washed three times with PBS containing 0.075% Tween-20 (PBST) and blocked by 2% bovine serum albumin (BSA) dissolved in PBST at 30 ℃ for 1 hour. For detection of Gal d2-specific IgG, IgG1 and IgG2a, sera samples serially diluted at two-fold (starting from 1: 400) were added into the wells and incubated for 2 hours at 30 ℃. Then HRP-conjugated rabbit anti-mouse IgG antibodies (1: 50,000 dilution, Abcam)，IgG1 antibodies (1: 5000 dilution, Southern Biotech) or IgG2a antibodies (1: 5000 dilution, Southern Biotech) were added and incubated for 1 h at 30 ℃, respectively. For detection of Gal d2-specific IgE titer, sera samples (1: 10 diluted) were added to the wells and incubated for 2 hours at 30 ℃, followed by incubation with HRP-conjugated rabbit anti-mouse IgE antibodies (Southern Biotech, 1: 5000 dilution) for 1 hour at 30 ℃. After washing steps, TMB substrate was used for development and the absorbance was measured at 450 nm using the SpectraMax^®^ Absorbance Reader (Molecular Devices).

**Enzyme Linked Immunosorbent Assay (ELISA)**

Level of serum IFN-γ or IL-10 in the culture supernatants were measured using commercial ELISA kits purchased from MultiSciences Biotech and Elabscience Biotec, respectively. Measurements were performed according to the manuals. Tecan sunrise Microplate reader was used for the detection of the absorbance at 450 nm.

**Competitive ELISA assay**

To assess the allergen-blocking capacity of vaccine-induced antibody, sera samples collected from mice that had been immunized with three doses of mRNA vaccines at an interval of 7 days were used. 96-well plates were pre-coated with 100 µg/mL Gal d2 and incubated overnight at 4 ℃. Upon washing with PBST and blocked by 2% BSA at 37 ℃ for 2 h, sera collected from unvaccinated naive mice or vaccinated mice (n=5) were heat-inactivated at 56 ℃ for 2 hours and then added into the plates for incubation at 4 ℃ for 1 hour. PBS was added into additional wells and served as negative control. Following washing steps, sera samples containing Gal d2-specific IgE that was collected from Gal d2-allergic mice were 1: 10 diluted and added to the wells for overnight incubation at 4 ℃. After washing, HRP-conjugated rabbit anti-mouse IgE antibodies (1:5000 dilution) were added for incubation at 30 ℃ for 1 hour. After washing, TMB substrate was used for development, and the absorbance was measured at 450 nm. The inhibition rate (%) was calculated as follows: 100 − [(OD_i_ / OD_PBS_) × 100], where OD_i_ represents the optical density value of test mouse sera from mRNA vaccine immunized mice or from naïve mice, and OD_PBS_ denotes the optical density of the wells with PBS added.

**Enzyme-Linked Immunospot (ELISPOT) Assay**

Frequencies of Gal d2-specific IL-2, IFN-γ, or IL-10 secreting T cells were measured using commercial ELISpot kits purchased from MABTECH according to the instruction. In brief, murine splenocytes (0.2 million cells per well) were incubated with or without overlapping peptide pool (Miltenyi) at the concentration of 10 μg/mL for 24 hours. Cells stimulated with S. aureus Enterotoxin Type B Toxoid (SEB, Creative Diagnostics) were treated as positive control. The primary antibody (detection antibody coupling biotin) was diluted with antibody diluent at the ratio of 1:1000 (final concentration: 1μg/mL) and added to each well and plates were incubated for 2 hours at room temperature. Following washing steps, secondary antibody (Streptavidin-ALP) was diluted with antibody diluent at 1:1000 ratio and added to each well, and plates were then incubated for 1 hour at room temperature. After washing, filtered BCIP/NBT-plus solution was added to each well for color development in dark. Spots were counted using CTL-Immunospot S6 Analyzer. Results were depicted as spot-forming cell (SFC) per million stimulated cells.

**Evaluation of basophil activation**

Frequencies of activated basophils (identified as IgE^+^CD200R3^+^CD63^+^ cells) were quantified using flow cytometric assay. In brief, 2×10^6^ murine splenocytes were washed with PBS and stained with LIVE/DEAD™ Fixable Aqua Dead Cell Stain Kit (Thermo) for 5 minutes and then incubated with antibody cocktails and Fc receptor blocking reagent (Miltenyi) for 20 minutes at 4°C in dark. Antibodies used in this analysis include anti-mouse CD45-Pacific Blue™ (clone: 30-F11, Biolegend), anti-mouse IgE-PE (clone: RME-1, Biolegend), anti-mouse CD49b-Percp-Cy5.5 (clone: HMα2, Biolegend), anti-mouse CD200R3-AF647 (clone: Ba13, Invitrogen), anti-mouse CD63-AF700 (clone: NVG-2, Biolegend). Flow cytometric analysis was performed on BD FACSCelesta. Data were analyzed with FlowJo software (version 10.8.1).

**Statistical analysis**

Statistical calculations were performed using GraphPad Prism v8.0. Comparisons between two groups utilized the Mann-Whitney U test. Statistical difference among three different groups was analyzed by one-way ANOVA test. p value less than 0.05 was considered statistically significant (*p ≤ 0.05, **p ≤ 0.01, ***p ≤ 0.001, ****p ≤ 0.0001).
